# The trimeric thylakoidal Tat receptor complex consists of a homo-oligomeric TatC core with associated TatB subunits

**DOI:** 10.1101/2025.01.10.632409

**Authors:** Matthias Reimers, Mario Jakob, Ralf Bernd Klösgen

## Abstract

The Twin-arginine translocation (Tat) machinery, which is found in most cellular membranes containing a respiratory or photosynthetic electron transport chain, is characterized by its unique ability to catalyze membrane transport of folded proteins without impairing the membrane potential. In plant thylakoids, Tat machinery consists of three subunits, TatA, TatB, and TatC, with the latter two, TatB and TatC, forming membrane-integral multimeric TatBC receptor complexes. Here we have analyzed the stability and the subunit composition of these complexes after solubilization of thylakoids with the mild detergent digitonin as well as after additional affinity-purification. Employing different detergent combinations and/or heat treatment (40°C) followed by BN-PAGE and Western analysis we could identify four distinct Tat complexes with apparent molecular masses ranging from approximately 230 kDa to 620 kDa. Treatment of the largest Tat complex with either heat or detergents like DDM or Triton X-114 led to its stepwise breakdown into the three smaller complexes resulting from the successive release of TatB subunits from a relatively stable TatC core complex. From these data we conclude that the fully assembled, physiologically active TatBC receptor complex consists of a stable, trimeric TatC core to which three TatB subunits are bound independently from each other.

## Introduction

Among the mechanisms mediating protein transport across biological membranes the Twin-arginine translocation (Tat) pathway is particularly remarkable because it is able to transport proteins in a folded conformation. The Tat pathway is found in the majority of cellular membranes housing a respiratory or photosynthetic electron transport chain, i.e., in the cytoplasmic membranes of bacteria and archaea, in the thylakoid membranes of plants and cyanobacteria and even in the inner mitochondrial membranes of plants and basal eukaryotes [1–6]. Despite some differences in detail the overall structure of these Tat translocases and their presumed mechanism of operation seem to be conserved to a large extent.

Tat translocase is composed of the three integral proteins TatA, TatB, and TatC (in chloroplasts also called Tha4, Hcf106, and cpTatC, respectively). TatA and TatB are closely related bitopic membrane proteins with a single N-terminal transmembrane helix and a cytoplasmically (bacteria) or stromally (chloroplasts) exposed hydrophilic domain which consists of a short hinge region, an amphipathic helix and a less conserved soluble region [7–10]. TatC comprises six transmembrane helices, which are connected by small interhelical loops, and cytoplasmically/stromally exposed N- and C-terminal domains [11,12].

TatB and TatC constitute integral heteromeric complexes of approximately 600 kDa, which act as membrane receptors for precursor proteins carrying Tat-specific signal peptides [13–16]. In contrast, the role of TatA, which seems to be essential solely for the actual membrane translocation step [17–19], is still a matter of debate. Several working models are being discussed including (i) a multimeric TatA transmembrane pore [8,20–22], (ii) a membrane destabilizing function of TatA caused by its seemingly short N-terminal transmembrane helix [23–26], and (iii) a catalytic, co-enzymatic function of TatA activating the integral TatBC complex to also perform the actual membrane translocation of the Tat substrate [19].

Despite the principal agreement on the functional role of the TatBC receptor complexes, their structure is still not known. Upon BN-PAGE, digitonin-solubilized thylakoidal Tat receptor complexes of pea appear as one or two bands in the range of 560 - 620 kDa [13–16], which were shown to contain TatB and TatC in an approximately 1:1 stoichiometry [14,27,28]. While in bacteria TatA has repeatedly been found in association with such Tat receptor complexes [29–35], TatA could not be detected in the thylakoidal receptors [15,27]. Also, the oligomerization state of the Tat complexes remains controversial with models ranging from dimer to octamer, although trimer and tetramer models have recently been favored [14,15,36–43].

In this manuscript, we have used detergent and heat treatment of digitonin-solubilized thylakoids as well as of affinity-purified thylakoidal Tat receptor complexes in combination with Blue Native-PAGE and Western analyses to dissect these complexes in a controllable manner. With this, we show the relationship between complex size and the TatB-TatC ratio and take a new step in understanding the structure of this unique protein translocase.

## Materials and Methods

### Generation and purification of polyclonal antisera against TatB and TatC from *Pisum sativum*

Polyclonal antisera were raised by immunizing rabbits with the C-terminal stromal domain of pea TatB (residues 109 to 261, NCBI Refseq NP_001413888.1) or the N-terminal stromal domain of pea TatC (residues 53 to 136, NCBI Refseq NP_001413900.1) following the protocol of Green & Manson [44]. Both antigens were obtained by heterologous overexpression using the pET-system [48] and *E. coli* strain BL21(DE3) with vector pET-30a (+) (Novagen). Each antigen was then purified by immobilized metal ion affinity chromatography (IMAC) on HiTrap Chelating HP columns (GE Healthcare) according to the manufacturer’s instructions. The polyclonal antibodies obtained from immunization were affinity purified according to Narhi et al. [45], following the protocol of Zinecker et al. [46].

### Cloning of tagTP for heterologous expression or cell free translation

The cDNA fragment encoding the entire transit peptide of the OEC16 precursor protein from *Spinacia oleracea* (residues 1 to 83, NCBI Refseq NP_001413330.1) was cloned into the NcoI and XhoI restriction sites of plasmid pET30a (+). The resulting construct encodes a polypeptide comprising an N-terminal combined His-/S-peptide-tag in frame with the transit peptide. In order to increase the affinity of the Tat-specific signal peptide of OEC16 to the Tat receptor complexes, two modifications (S98C and M108F) were introduced by site-directed mutagenesis. The resulting construct, which encodes the Tat substrate analogon termed tagTP (Fig. 5A), was furthermore cloned into plasmid pBAT [47] to facilitate synthesis of the peptide also by *in vitro* transcription/translation.

### Overexpression and purification of tagTP

The Tat substrate analogon tagTP was obtained by overexpression in *E. coli* using the pET-system [48]. After induction for 3 hours, the cells were disrupted by French Press at 1000 psi, centrifuged and the pelleted inclusion bodies were dissolved in Ni^2+^-affinity column binding buffer (20 mM HEPES, 500 mM NaCl, 20 mM Imidazole, 6 M guanidine hydrochloride, pH 7.5). Insoluble material was removed by ultra-centrifugation at 140.000 × g and the supernatant was used for IMAC. After sample loading, the column was washed with 6 column volumes of binding buffer. TagTP was recovered in three steps using 0.5, 1 and 0.5 column volumes of elution buffer (20 mM HEPES, 500 mM NaCl, 500 mM Imidazole, 6 M guanidine hydrochloride, pH 7.5), respectively (Fig. S2).

### Binding of tagTP to thylakoidal TatBC complexes

TagTP was dialyzed against 10 mM HEPES, 5 mM MgCl2, 1 M urea, pH 8.0, using dialysis tubes with a molecular mass cut-off of 1 - 3.5 kDa. For Tat complex binding, 1 volume of this tagTP solution (approximately 300 μg/ml = 20 μmol/l) was incubated for 20 min on ice with 4 volumes freshly isolated pea thylakoids (750 μg chlorophyll/ml). Alternatively, 5 μl radiolabeled tagTP obtained by *in vitro* translation in the presence of ^35^S-methionine using the Flexi^®^ Rabbit Reticulocyte Lysate System (Promega) were incubated with 40 μl pea thylakoids (750 μg chlorophyll/ml). After incubation, the thylakoids were sedimented (5 min, 10.000 × g, 4°C), washed once with HM buffer and resuspended in 40 μl HM-buffer.

### Solubilization of thylakoidal membrane complexes

Pea thylakoids resuspended in HM buffer (750 μg chlorophyll/ml) were supplemented with 1 volume of two-fold concentrated solubilization buffer (3% digitonin, 10 mM Bistris, pH 7.0, 1 M NaCl, 20% glycerol, 1 mM EDTA, 0.1 mM MgCl2, 0.2 mM PMSF) resulting in a final detergent concentration of 1.5 % digitonin. After incubation for 1 h at 7°C, the assays were centrifuged twice (13.000 × g, 4°C, and 200.000 × g, 4°C) and the solubilized membranes were recovered with the supernatant.

### Affinity purification of TatBC complexes

2 ml thylakoid suspension preincubated with tagTP and solubilized with digitonin were supplemented with imidazole buffer (final concentration 20 mM) and subjected to IMAC on His Spintrap columns (GE Healthcare). The columns were washed twice with 600 μl each of the same buffer and eluted with 4 × 100 μl solubilization buffer supplemented with 500 mM imidazole. For mass spectrometry, the elution fractions were pooled, diluted with solubilization buffer to an imidazole concentration of 30 mM, and subjected once more to Ni^2+^-affinity chromatography as described above.

### Blue Native PAGE

Blue Native polyacrylamide electrophoresis (BN-PAGE) was carried out based on the work of Schägger et al. and Schägger & von Jagow [49,50]. The protocol was adapted for thylakoid samples according to Berghöfer et al. [13], except that digitonin was omitted from the mini gels used (10 × 10 × 0.1 cm). The Coomassie dye retained in the gels after electrophoresis did not interfere with the subsequent Western analysis and was therefore not removed.

### Western analysis

Proteins were transferred after electrophoresis to PVDF membranes using a semidry blotting system and buffers according to Kyhse-Andersen [51]. For the detection of thylakoidal TatC, it was crucial to omit the generally used treatment of the BN-gels with SDS-containing buffers (Fig. S3). After transfer, the membranes were washed with PBS buffer supplemented with 0.09 % hydrogen peroxide to suppress unspecific signals in the subsequent ECL reaction (Fig. S3).

### Miscellaneous

Isolation of chloroplasts and thylakoids from pea seedlings (*Pisum sativum* var. Feltham First) followed the protocol described by Hou et al. [52]. For autoradiography, gels were vacuum dried onto filter paper, exposed to phosphorimager screens and analyzed with a Fujifilm FLA-3000 (Fujifilm, Düsseldorf, Germany) using the software package BASReader (version 3.14). For mass spectrometry, the samples were prepared and analyzed by LC-MS/MS as described by Wittig et al. [53]. The resulting data were aligned against the sequences of tagTP, pea TatB, pea TatC and the proteome of *Cicer arietinum* (chickpea), a close relative of *Pisum sativum*.

## Results

### Solubilized Tat complexes can be stepwise converted into three smaller TatC-containing complexes

A suitable method for the analysis of membrane complexes is the solubilization of the respective membranes with detergents like digitonin followed by Blue Native polyacrylamide gel electrophoresis (BN-PAGE) and Western analysis. Employing anti-TatC antibodies, we could identify with this approach a complex of approx. 620 kDa as the predominant form of the presumed Tat receptor (Fig. 1, complex #1) and a smaller complex of approx. 560 kDa (#2), which was usually found to a somewhat lower extent. In some instances, an even smaller complex of approx. 440 kDa (#3) could be identified, though with usually only low signal intensity. Complexes #1 and #2 are reminiscent in size to those Tat receptor complexes observed by indirect labelling with the radiolabeled Tat substrate 16/23 [13].

**Figure 1:**
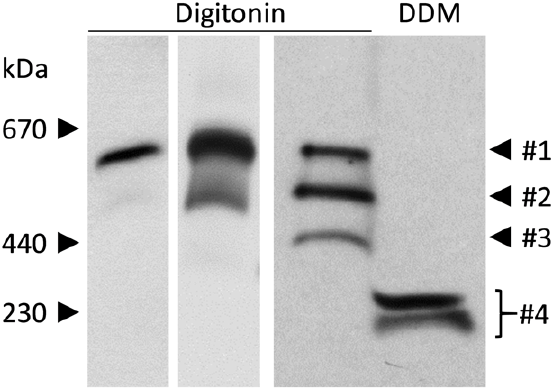
Variability in the electrophoretic patterns of thylakoidal Tat complexes. Thylakoid membranes corresponding to 7.5 μg chlorophyll were solubilized for 1 h at 4 °C with either 1.5% *digitonin* or 1.5% *DDM*. After centrifugation for 20 min at 140,000 × g, the supernatants were subjected to electrophoresis on a 5–13.5% detergent-free Blue Native polyacrylamide gradient gel. After Western transfer to PVDF membrane, TatC-containing protein complexes were decorated with affinity-purified antisera raised against the stromal domain of TatC from pea. The size of marker protein complexes is shown on the left side, while the four TatC-containing complexes observed (#1 - #4) are indicated on the right. For thylakoids solubilized with *digitonin*, the results of three independent experiments are shown.

In contrast, solubilization of the thylakoids with DDM instead of digitonin showed a TatC-containing complex with an apparent molecular mass of only approximately 230 kDa, which sometimes appears as a double band on such gels (Fig. 1, #4). Remarkably, when digitonin-solubilized thylakoid samples were additionally supplemented with rising concentrations of DDM (0.07% - 1.5%), Tat complex #1 was gradually converted into complexes #2, #3 and #4. Complex #4 appears to be a kind of stable core complex as it could not be further degraded by further increase in the DDM concentration (Fig. 2).

**Figure 2:**
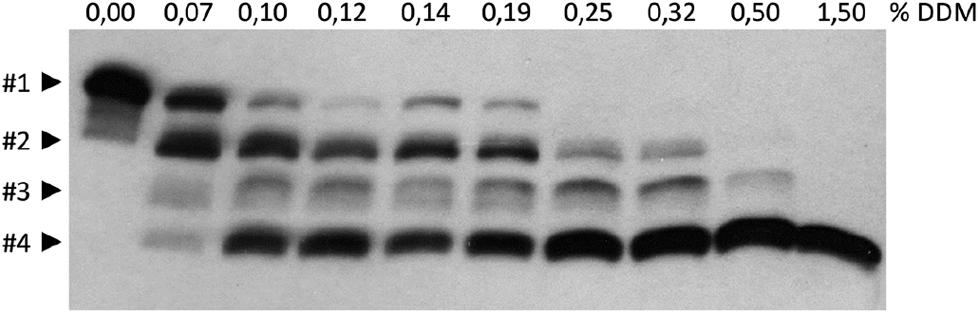
Increasing concentration of DDM leads to stepwise size-reduction of digitonin-solubilized Tat complexes. Digitonin-solubilized pea thylakoids were treated with increasing amounts of *DDM*, incubated for 5 min at 4°C and analyzed by BN-PAGE and Western analysis employing anti-TatC antibodies. The final DDM concentration in each sample (in *per cent*) is given above the lanes. For further details see the legend to Fig. 1.

Essentially the same result was obtained when DDM was replaced by Triton X-114 in such experiments (Fig. 3). Again, at low concentration of Triton X-114 (≤ 0.2%), digitonin-solubilized Tat complex #1 was converted into the smaller complexes #2, #3 and #4, while at high Triton X-114 concentration (≥ 0.5%) predominantly complex #4 was found. It should be noted that the apparent size of the complexes on the gel appears to vary to some extent depending on the type and concentration of the detergent used. Still, the results obtained with both, DDM and Triton X-114, are strikingly similar with respect to the ladder-like pattern and the size of the terminal complex #4, and point to a stepwise release of peripheral components from a TatC-containing core complex which takes place independently of the detergent used.

**Figure 3:**
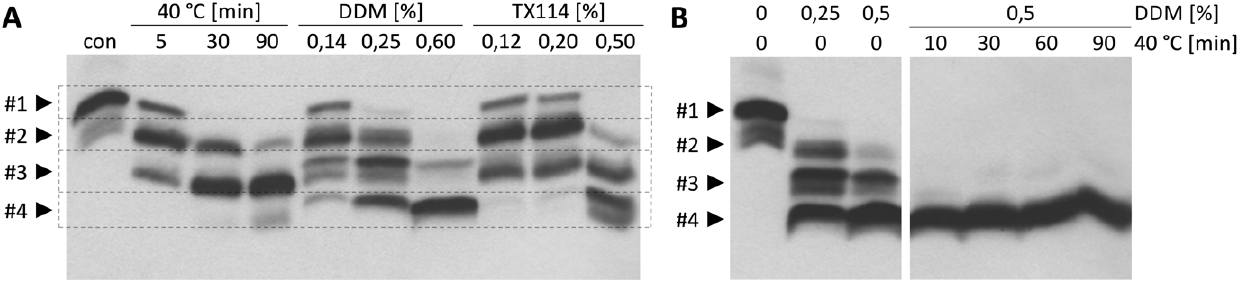
DDM, Triton X-114 and heat treatment have similar effects on the stability of digitonin-solubilized Tat complexes. **(A)** Digitonin-solubilized pea thylakoids (*con*) were incubated for *5, 30* or *90 min* at *40 °C*, or treated with increasing amounts of either *DDM* (*0*.*14 - 0*.*60%*) or Triton X-114 (*TX114, 0*.*12 - 0*.*50%*), as indicated above the lanes, and analyzed as described in the legend to Fig. 2. (**B)** Digitonin-solubilized pea thylakoids were treated with either *DDM* (*left panel*) or simultaneously with *DDM* and *40°C* (*right panel*) as indicated above the lanes. See text for further details.

Even heat treatment of digitonin-solubilized thylakoids had principally the same effect (Fig. 3 A). When such samples were incubated for 5, 30, or 90 min at 40° C, Tat complex #1 was successively converted into complexes #2 and #3 over time. After 90 minutes, also complex # 4 was found, though at only minor amounts, which might be indicative of higher stability of complex #3 in the absence of additional detergent.

Finally, if digitonin-solubilized thylakoids were treated with a combination of DDM plus heat prior to analysis, exclusively complex #4 was found (Fig. 3 B). Neither of the larger complexes #1, #2, or #3 nor any structure smaller than complex #4 could be detected, which is in line with the assumption that this complex represents a minimal, stable Tat core complex comprising TatC.

### The conversion of Tat complexes into smaller forms correlates with a dissociation of TatB

While the analysis so far has focused exclusively on TatC, we wanted to analyze also TatB which is assumed to be present in the Tat complexes in equimolar amounts to TatC. In fact, Western analyses after BN-PAGE of digitonin-solubilized thylakoids showed identical signals corresponding to complexes #1 and #2 when developed with antibodies raised against either TatB or TatC (Fig. 4, lane 0 min). The only difference is an additional TatB-specific signal in the lower part of the gel (open arrowhead) which presumably represents non-assembled TatB subunits. When such digitonin-solubilized thylakoids were additionally treated with either DDM, Triton X-114, or heat (40°C), the results obtained for TatB were largely similar to those of TatC, as exemplarily shown in Fig. 4 for heat treatment. However, there was one significant difference: while complexes #1, #2 and #3 were recognized by both types of antibodies, complex #4 could exclusively be detected by anti-TatC but not by anti-TatB antibodies, not even after overexposure of the membrane (Fig. 4, right panel). Complex #4 therefore does not appear to contain TatB protein. From these observations we conclude that the decrease in size of the Tat complexes #1 to #4 corresponds to a stepwise release of TatB subunits and that complex #4 represents a relatively stable TatC core complex that is devoid of any TatB subunit.

**Figure 4:**
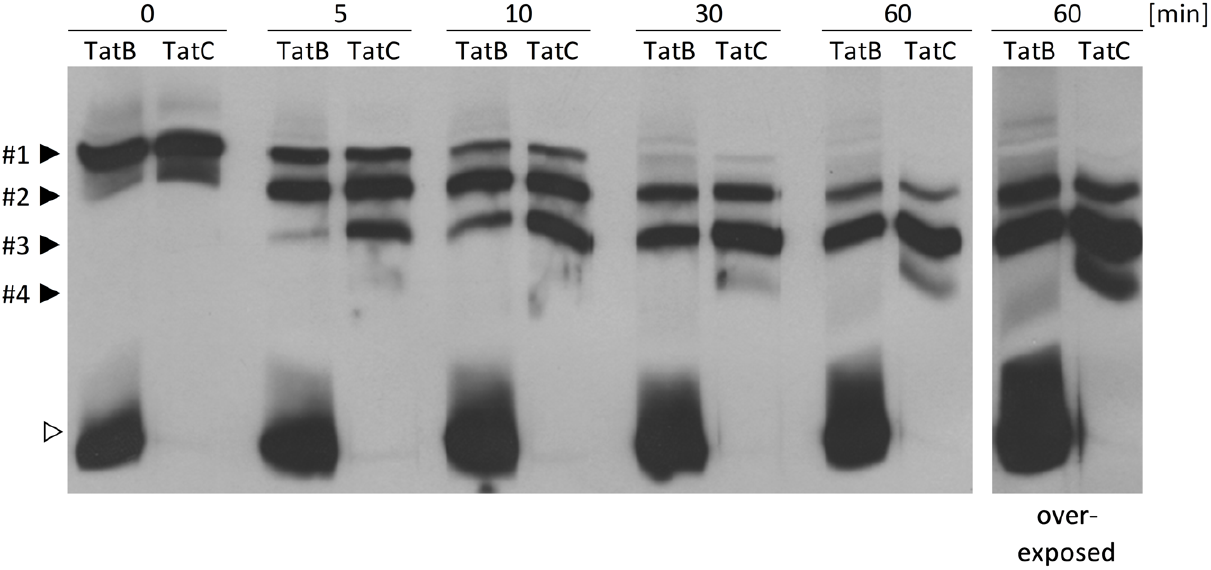
Heat-induced size reduction of solubilized Tat complexes results from stepwise release of TatB subunits. Digitonin-solubilized thylakoids were incubated at 40 °C for time periods ranging from *0 - 60 min* as indicated above the lanes. After BN-PAGE and Western transfer to PVDF membrane, each lane was cut longitudinally and the two halves were decorated separately with affinity-purified antisera raised against either TatB or TatC. Prior to ECL development, the corresponding halves were reassembled to allow for the immediate comparison of the results (Fig. S1). Exposition time: 1 min (overexposed: 2 min). The open arrowhead indicates a low molecular TatB species (see text). For further details see the legend to Fig. 1.

**Figure 5:**
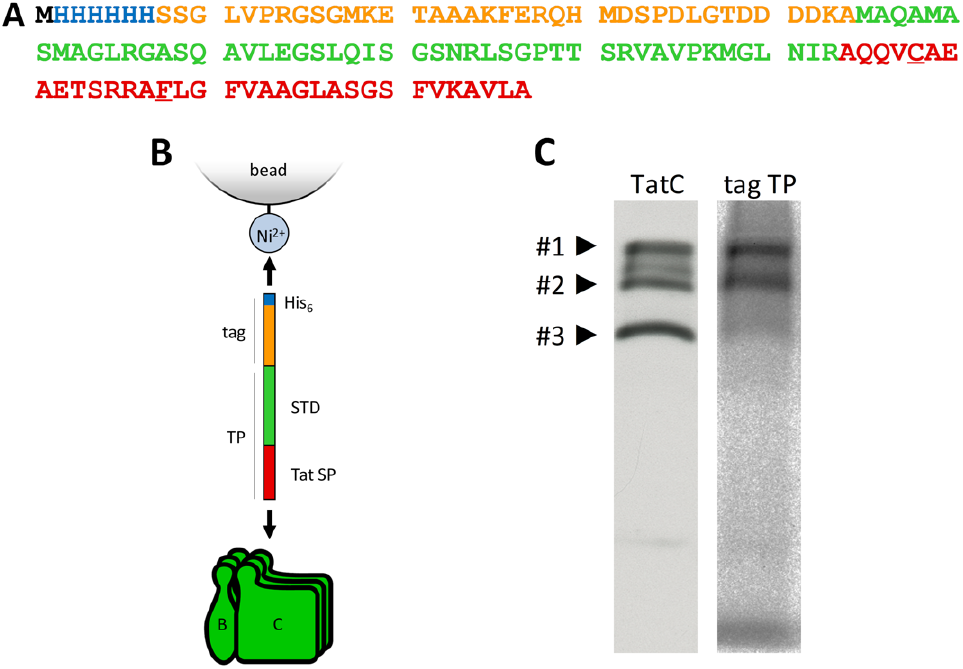
The Tat substrate analogon tagTP binds specifically to the thylakoidal TatBC receptor complexes. **(A)** Amino acid sequence of the Tat substrate analogon tagTP given in the one-letter-code. The colors of the fonts represent different parts of the construct (blue - N-terminal His6-tag, orange - S-peptide-tag, green - stroma-targeting domain of the spinach OEC16 precursor, red - signal peptide/thylakoid-targeting domain of the spinach OEC16 precursor). The two amino acid residues modified for increased Tat receptor affinity (S98C and M108F) are *underlined* (see Materials and Methods for further details). Schematic representation of the interaction of tagTP with both, IMAC beads and the Tat receptor complex. (**C)** Isolated pea thylakoids were incubated for 20 min on ice with radiolabeled tagTP obtained by *in vitro* translation in the presence of [^35^S]-methionine. After solubilization with digitonin, the thylakoids were separated by BN-PAGE followed by either Western analysis employing anti-TatC antibodies (left panel) or by autoradiography (right panel). For further details see the legend to Fig. 1.

In line with this assumption, the amount of presumably non-assembled TatB appears to increase with increasing time of heat treatment (Fig. 4) which might reflect the accumulation of TatB subunits released from the Tat complexes upon heat treatment. However, the large amount of such TatB proteins even in non-treated digitonin-solubilized thylakoids prevents any firm conclusion, which is why we next tried to purify the Tat complexes from the thylakoid membrane.

### Affinity-purification of Tat complexes from thylakoid membranes

To achieve such a purification and to also compensate for the low abundance of the Tat complexes in the thylakoid membrane, we aimed to enrich these complexes by immobilized metal affinity chromatography (IMAC). For this purpose, we took advantage of the receptor function of the Tat complexes, i.e., of their property to bind substrate proteins carrying Tat-specific signal peptides (e.g. [13]). As a bait peptide, we generated a His6-tagged variant of the transit peptide of the 16 kDa subunit of the oxygen evolving complex (OEC16) which contains a Tat-specific signal peptide that was further optimized for binding to the Tat receptor complex (for details see Materials and Methods). The stroma targeting domain of the transit peptide, which does not impair binding of the Tat signal peptide to the Tat receptor [13], was retained as spacer to the N-terminal His6-tag region (Fig. 5 A and B).

Functional binding of this Tat substrate analogon named tagTP (tagged transit peptide) to the Tat complexes was demonstrated by incubating a radiolabeled version of it with isolated thylakoids. The thylakoids were then solubilized with digitonin, separated by BN-PAGE and finally analyzed by both, autoradiography and Western detection of TatC. As shown in Fig. 5C, the autoradiography signals obtained with tagTP comigrate with the TatC containing complexes #1 and #2 demonstrating that this Tat substrate analogon can efficiently bind to both receptor complexes. In contrast, complex #3 seems to bind only marginal amounts of tagTP despite the fact that it is clearly visible upon Western detection of TatC.

For the pursued enrichment of the Tat complexes, micromolar quantities of tagTP were produced by heterologous overexpression in *E. coli* followed by purification using IMAC (Fig. S2). After incubation of such purified tagTP with isolated thylakoids and subsequent solubilization with digitonin, the bait peptides together with potentially bound Tat complexes were recovered by IMAC. All samples were subsequently analyzed by BN-PAGE followed by Western detection of TatC to identify TatC-containing membrane complexes. As shown in Fig. 6A, the flow through of the IMAC-column lacks essentially all TatC-containing complexes, suggesting that the majority of these complexes had bound tagTP and were thus retained by the affinity matrix. Indeed, the Tat complexes were strongly enriched in elution fractions 1 - 4. In line with the results shown in Fig. 5C, predominantly complexes #1 and #2 were found, while only traces of complex #3 and no complex #4 could be detected in the elution fractions.

**Figure 6:**
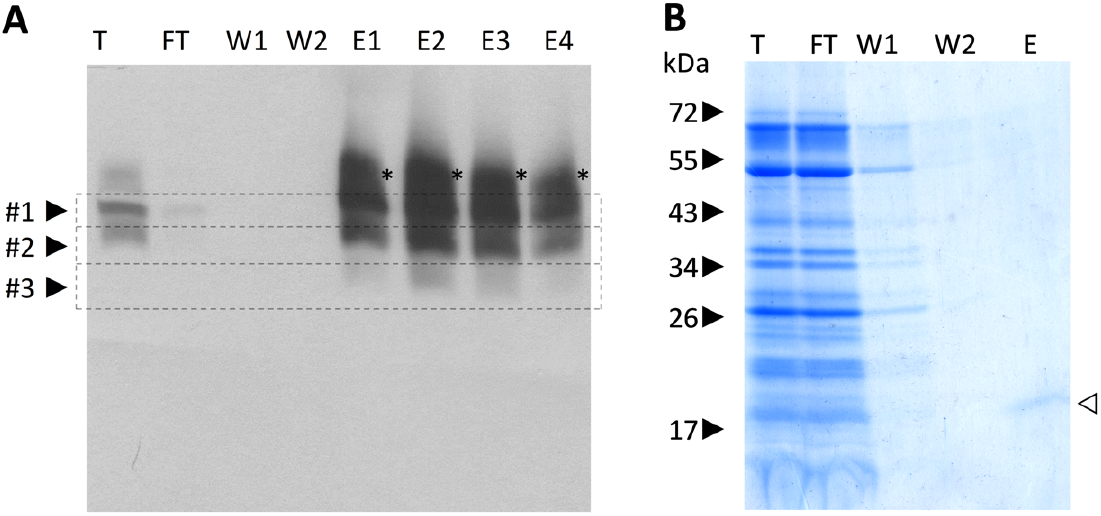
Affinity-purification of thylakoidal TatBC complexes using tagTP as a bait. Isolated pea thylakoids were incubated for 20 min on ice with affinity-purified tagTP obtained by overexpression in *E. coli*. After washing and solubilization with digitonin, the sample was loaded onto an IMAC column. Stoichiometric amounts of loading sample (T), flow through (FT) and wash fractions (W1, W2) were loaded, while the elution fractions (E1-E4) were approximately 3.5-fold concentrated. The samples were subjected to **(A)** BN-PAGE followed by Western analysis with anti-TatC antisera or **(B)** SDS-PAGE and subsequent Coomassie Colloidal staining. The asterisk points to an additional signal which was specifically observed only after incubation with tagTP optimized for Tat complex binding (see text for details). In (**B)**, an aliquot of the pooled elution fractions 1 - 4 was loaded. The open arrowhead indicates the position of tagTP. For further details see the legend to Fig. 1.

It should be noted that, in some of these experiments, an additional signal at even higher molecular mass than complex #1 was observed by both, TatB and TatC detection (marked with an asterisk in Fig. 6A). Although the nature of this signal is yet elusive, its appearance seems to correlate with the M108F replacement in tagTP (Fig. S4), in line with published data analyzing an identically modified maize homologue of pea OEC16 [54].

In order to assess the quality of the affinity purification procedure, all samples were additionally analyzed by SDS-PAGE and Coomassie staining. As shown in Fig. 6B, the flow through of the IMAC-column contained virtually all proteins that were present also in the loading control (lane T), in line with the presumed selectivity of the affinity column. In contrast, in the pooled elution fractions (lane E) solely tagTP could be detected but neither TatB nor TatC nor any other protein. This was rather unexpected in view of the strong Western signal obtained for TatC after BN-PAGE analysis (Fig. 6A). At first glance, this might suggest that each Tat complex had bound excess amounts of tagTP which would consequently be easily detectable. However, we prefer an alternative explanation instead: it has repeatedly been shown that Tat substrates like the chimeric 16/23 protein are capable of binding not only to Tat receptor complexes but that they have an intrinsic property to insert into lipid membranes independently of any proteinaceous transport machinery [52,56–58]. It therefore appears possible that also the Tat substrate analogon tagTP is able to bind not only to Tat complexes but to insert also directly into the lipid bilayer of the thylakoid membrane. As a result, such thylakoids would not only contain tagTP bound to the Tat receptor but also unpredictable amounts of free tagTP in the membrane, which likewise would be recovered by IMAC.

This assumption is supported by the results of mass spectrometry analyses which were performed to determine the content of the purified Tat complex samples in more detail. For this purpose, the affinity purification of the Tat complexes was repeated, including two successive cycles of IMAC in order to further improve the degree of purity. The final elution fraction was subjected to SDS-PAGE and the whole gel lane was cut into pieces that were treated with trypsin. The released peptides were analyzed by LC-MS/MS and the results for all gel pieces were combined. The most prominent polypeptide that could be identified was tagTP, followed by matches for TatC and TatB at rank 3 and 4, respectively (Fig. S5). At rank 2, a polypeptide with an alignment score of only −2 is listed that did not match with any entry in the database. Subunits of the ubiquitous photosystems, like PsaN, PsaF or PsbS, were found to a clearly lesser extent than TatB and TatC despite the fact that they are highly abundant in the thylakoid membrane, which confirms the selectivity and suitability of the chosen purification procedure. It is particularly remarkable though, that TatA, the third component of Tat translocase, could not be detected at all in the samples which clearly demonstrates that, in thylakoids, TatA is not part of the Tat receptor complex, at least not at the stage of precursor binding, before the actual membrane translocation process has started.

### Tat complexes have a trimeric architecture and consist of a TatC oligomer with less stably associated TatB

Finally, we subjected the tagged and affinity-purified Tat complexes also to the treatments with detergents, like DDM or Triton X114, and/or heat in the same way as described above. In all these experiments, essentially the same results were obtained as with the solubilized thylakoids. This is exemplarily shown in Fig. 7 which illustrates the stepwise decrease in size of the purified Tat complex after DDM-treatment. As shown by Western analysis with both, anti-TatC and anti-TatB, such a treatment ultimately gives rise to a TatC core complex (#4) which is devoid of any TatB. Here, unlike the results shown in Fig. 4, untreated samples did not display any TatB-specific signals in the lower sections of the gel and, therefore, the accumulation of non-assembled TatB molecules upon DDM-treatment is clearly visible (open arrowhead, Fig. 7). As suggested already above, this result now clearly demonstrates that the reduction in complex size observed under these experimental conditions is indeed correlated with the release of TatB subunits from the TatBC receptor complexes #1 - 3.

**Figure 7:**
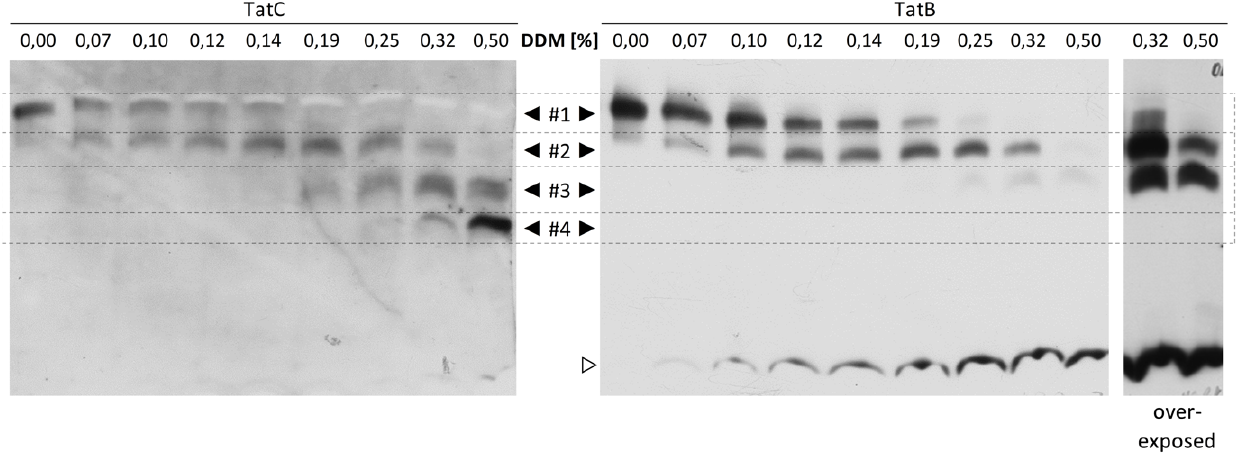
Stepwise release of TatB from Tat receptor complexes gives rise to a homo-oligomeric TatC core complex. Digitonin-solubilized and affinity-purified Tat complexes were treated with increasing amounts of DDM, incubated for 5 min at 4°C and analyzed by BN-PAGE and Western analysis employing either anti-TatC (left panel) or anti-TatB antisera (right panel). The final DDM concentration in each assay (in per cent) is given above the lanes. Exposition times: anti-TatC - 1 min, anti-TatB - 30 s or 180 sec (overexposed). For further details see the legends to Figs. 1 and 6.

Considering the number of Tat complexes of intermediate size observed, we furthermore conclude that the thylakoidal Tat receptor complex has a trimeric architecture consisting of a homo-oligomeric TatC core with less firmly associated TatB subunits (Fig. 8).

**Figure 8:**
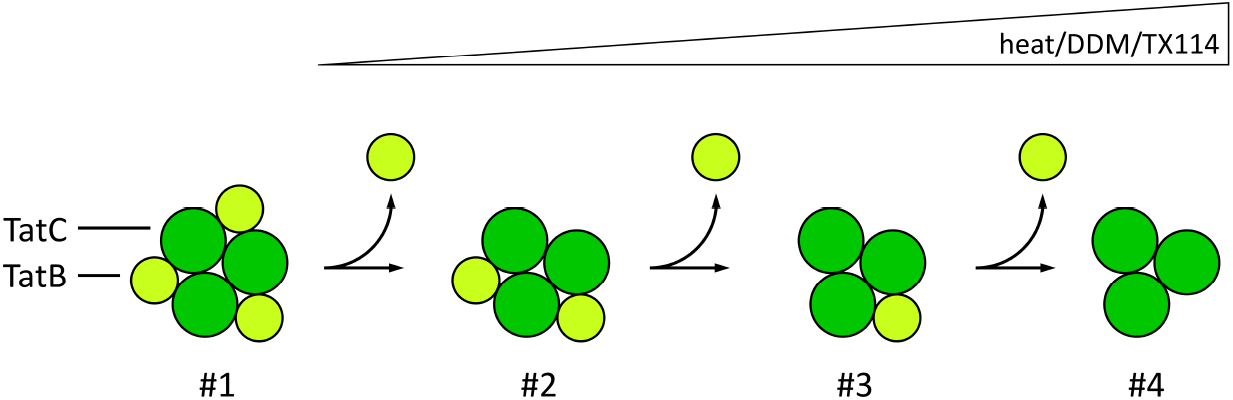
Model of Tat complex dissociation. Illustration of the proposed loss of TatB subunits from the proposed trimeric TatBC receptor complex. Upon treatment with heat, DDM or Triton X-114, digitonin-solubilized TatB3C3 complex #1 stepwise releases single TatB subunits leading to the Tat complexes #2 (TatB2C3), #3 (TatB1C3) and, finally, the homo-oligomeric TatC core complex #4 (TatC3).

## Discussion

It was the goal of this work to gain insight into the structure and composition of the plant thylakoidal Tat receptor complex. Our results indicate that the active receptor complex has a trimeric architecture which consists of a relatively stable core complex composed exclusively of TatC subunits to which equimolar amounts of TatB subunits are less tightly bound.

### TatBC complexes consist of a stable TatC core and loosely attached TatB subunits

After solubilization of thylakoidal membrane complexes with digitonin, a Tat complex with a molecular mass of approximately 620 kDa (#1) can be identified by both, substrate labeling and Western analysis. This complex is assumed to be the physiologically active receptor of thylakoidal Tat translocase which is capable of binding Tat substrates carrying a Tat-specific signal peptide [13,27]. It can be successively converted into smaller complexes of approximately 560 kDa (#2), 440 kDa (#3) and 230 kDa (#4) by different treatments including heat (40°C) or additional detergent, like DDM or Triton X-114 (Figs. 2, 3, 4), which all lead to the release of TatB subunits. In some experiments, complex #2 even formed spontaneously upon solubilization with digitonin (e.g., Fig. 1), underlining the relative weakness of TatB association. Remarkably, such conversion to smaller complexes can be observed regardless of whether the treatment is performed with solubilized thylakoids or affinity-enriched Tat complexes (e.g., Figs. 2 and 7). Since complex #4, the putative TatC homo-oligomer, is quite stable and persists even under circumstances where all associated TatB is lost, we assume that TatC forms a robust scaffolding core complex to which TatB is less tightly bound.

When affinity-purified Tat complexes were used in such analyses, the TatB subunits released from the Tat complexes accumulated in non-assembled form of low molecular mass (Fig. 7). Intriguingly, even merely digitonin-solubilized thylakoids already contain a certain amount of this TatB moiety (Fig. 4), which might either represent a pool of unassembled TatB molecules or result from TatB release occurring too quickly after solubilization to be captured. Interestingly, pools of free TatB, but not of TatC, have been reported already earlier [15,27,59].

The idea of a relatively loose or dynamic TatB association is supported by results described by Zinecker et al. [46] who functionally replaced TatB in thylakoidal Tat complexes. In these experiments, the transport activity of Tat translocase in intact thylakoids was inhibited by affinity-purified anti-TatB antibodies. After supplementation of the assays with purified soluble TatB, transport activity could be restored which strongly suggests that in these assays an exchange of the inactivated TatB subunits present in the complexes with those added externally had taken place.

### The Tat receptor complex has a trimeric architecture

Based on the results shown in this manuscript, we propose a trimeric architecture of the thylakoidal Tat receptor complex (Fig. 8). We have deduced this structure simply by counting the number of observed complexes of intermediate size. Assuming that each of the three dissociation steps releases equal amounts of TatB from the “complete” Tat complex #1, which presumably has an equimolar stoichiometry of TatB and TatC subunits [14,28], the structure of this complex seems most likely to be a TatBC trimer composed of three TatB and TatC monomers each (leading to TatB3C3), although a trimeric structure of TatB and TatC dimers (leading to TatB6C6) cannot be strictly ruled out at present. Even bigger trimeric complexes (e.g., TatB9C9) appear highly unlikely considering the current literature data [14,15,36–43].

However, in neither of the two scenarios the molecular masses of the Tat complexes observed upon BN-PAGE correspond to those calculated from the molecular masses of the TatB and TatC subunits. While in native gels the Tat complexes showed apparent molecular masses ranging from 230 kDa (complex #4) to 620 kDa (complex #1), the calculated masses of these complexes are only between 100 kDa (complex #4) and 157 kDa (complex #1) if a TatB3C3 architecture is assumed. Even in a hypothetical TatB6C6 scenario, the calculated molecular masses of the four complexes would range only from 200 kDa to 313 kDa.

Of course, it is possible that additional constituents of the complex, which might have been missed to date, could contribute to the observed electrophoretic migration behavior. One obvious candidate is TatA, the third proteinaceous Tat component. However, we have so far never detected any TatA in thylakoidal Tat complexes, neither by mass spectrometry of the affinity-purified Tat complexes (Fig. S5) nor by any other analysis in our lab. In bacteria, on the other hand, TatA has been described to be present in Tat complexes [29–35]. Yet, even if stoichiometric amounts of TatA were present also in thylakoidal Tat complexes, it would still by far not be sufficient to explain the differences between the calculated and the observed molecular masses.

Instead, such deviation might be a consequence of the experimental approach used. For example, it is well known that both Coomassie and membrane lipids can affect the apparent molecular masses during BN-PAGE analysis [60–63]. In line with that, purified recombinant *E. coli* Tat complexes solubilized with the detergent glyco-diosgenin, which is structurally similar to digitonin, were found to contain large amounts of phospholipids [64]. Alternatively, Tat complexes might intrinsically exhibit an aberrant electrophoretic migration behavior. It was reported that even upon denaturing electrophoresis both, wild-type and mutant derivatives of TatB, showed a mobility that led to marked overestimation of its molecular mass [27,65,66]. It appears reasonable to assume that such overestimation might also hold true for BN-PAGE analyses.

Finally, comparing the TatB3C3 and TatB6C6 models with each other, the latter seems less likely because it would require at each dissociation step the coordinated release of two TatB molecules from the Tat complexes. Taking into account that there is no evidence for the presence of TatB dimers in such complexes, such a scenario is difficult to imagine. Hence, following the principles of Ockham’s razor we suggest the trimeric TatB3C3 architecture as the most likely structure of the thylakoidal Tat receptor complex.

## Supporting information

Graphical Abstract

## The abbreviations used are

Tat: twin-arginine translocation
OEC16: 16 kDa subunit of the oxygen-evolving system associated with photosystem II
Hepes: 4-(2-Hydroxyethyl)piperazine-1-ethanesulfonic acid
DDM: N-Dodecyl-β-D-maltoside
BN-PAGE: Blue native polyacrylamide gel electrophoresis
PMSF: phenylmethanesulphonyl fluoride
PVDF: polyvinylidene fluoride
IMAC: immobilized metal affinity chromatography

## Acknowledgements

We would like to thank Julian Bender and Carla Schmidt (current address: University of Mainz, Germany) for conducting the mass spectrometry analyses. Their contribution was essential to this study.

