## Supplementary Figures for "The trimeric thylakoidal Tat receptor complex consists of a homo-oligomeric TatC core with associated TatB subunits"

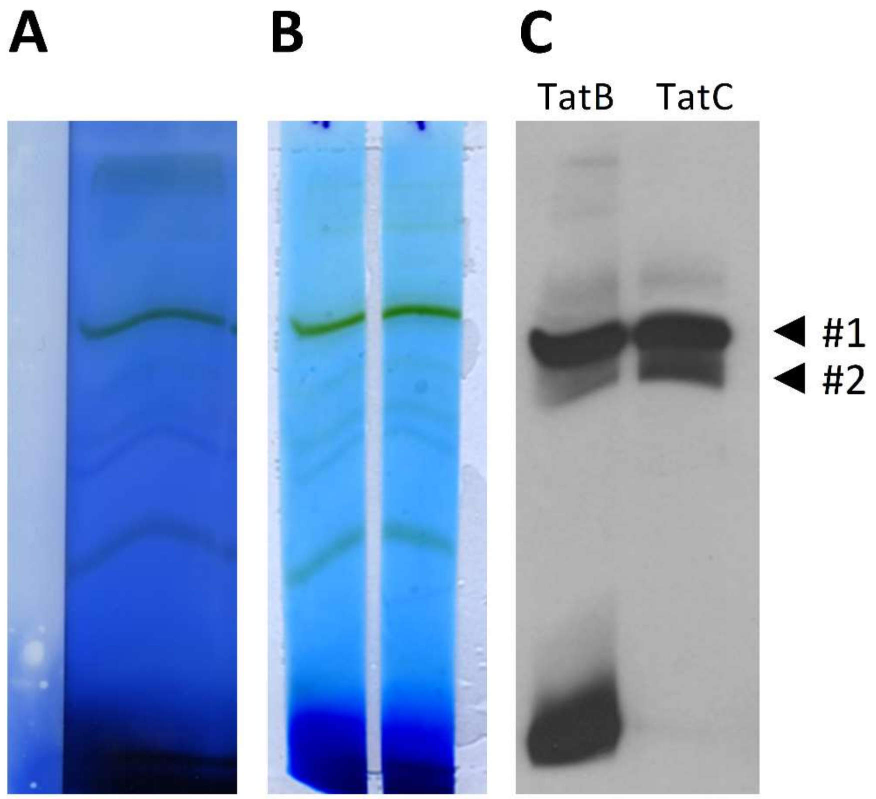


**Supplementary Figure S1: Working steps for Figure 4**

**(A)** Digitonin-solublized pea thylakoids were subjected to BN-PAGE with broad gel pockets that could accommodate solubilized thylakoids corresponding to 15 µg chlorophyll. A single lane (time point "0") is shown. After electrophoresis, the proteins were transferred to PVDF membrane and each lane was longitudinally cut. One half was incubated with *anti-TatB* antibodies, the other half with *anti-TatC* antibodies. **(B)** The halves were then reassembled leaving a small space in between and subjected to ECL detection **(C)**. The position of complexes #1 and #2 are indicated.

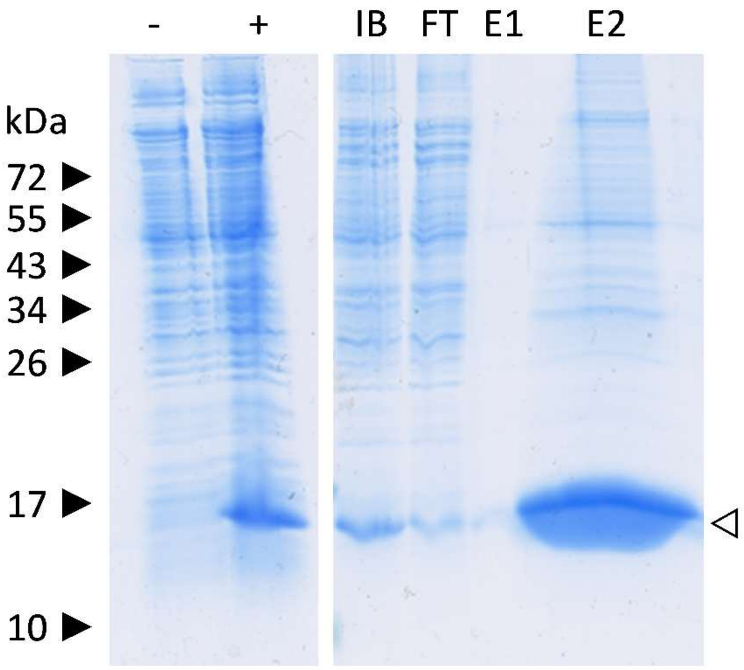


**Supplementary Figure S2: Heterologous overexpression and purification of tagTP**

The tagTP construct was overexpressed in *E. coli*. Whole cell extracts before (*lane* –) and after induction of expression (*lane +*) were analyzed by SDS-PAGE and Coomassie-Colloidal-Staining. After induction, the inclusion body fraction (*IB*), which contains most of the tagTP produced, was recovered, and subjected to IMAC. The flow through (FT) and elution fractions 1 (E1) and 2 (E2) are shown. Samples were analysed with SDS-PAGE and Coomassie staining.

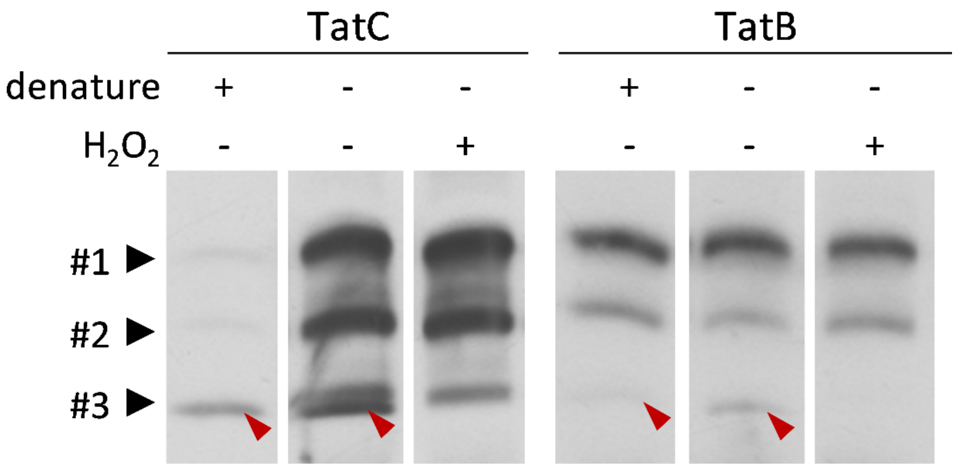


**Supplementary Figure S3: Critical optimization of BN-PAGE Western blotting procedure for analysis of thylakoidal TatC**

Digitonin solubilized pea thylakoids corresponding to 7.5 µg chlorophyll were separated by BN-PAGE on a 5 - 13.5 % PAA gel devoid of additional detergent. Subsequently, the gel was cut longitudinally along the borders of the lanes and the individual gel strips were prepared for Western Blotting with or without an additional denaturing step (22.5 mM Tris, 172.8 mM Glycine, 1.1 % SDS, 0.1 % Beta-Mercaptoethanol, 40 °C, 10 min). After the protein transfer, the PVDF membrane was also cut along the lane borders and selected strips were incubated with 0.09 % $H_2O_2$. The remaining working steps followed the Western Blotting protocol.

We found that it is critical to not denature BN-PAGE gels prior to protein transfer if TatC is to be analyzed, as this will greatly reduce the signal intensity up to complete absence of signals. This does not affect detection of TatB. We also found that treatment of Digitonin-solubilized thylakoids with Triton X-100 or X-114 led to a severe reduction or even total absence of TatC signals in such experiments. Also, $H_2O_2$ treatment of the blotted membranes removes an unspecific signal (red arrowheads) close to Tat complex #3 which presumably represents a reaction of the ECL solution with the cytochrome $b_6/f$ complex.

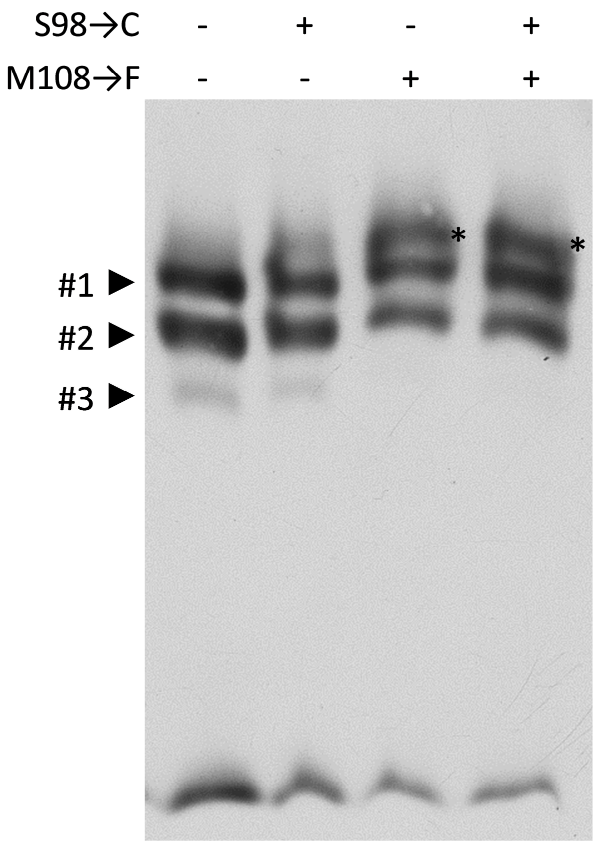


**Supplementary Figure S4: The M64F exchange in tagTP leads to the appearance of an additional high molecular weight signal upon BN-PAGE**

Affinity purification of thylakoidal Tat complexes using mutant derivatives of the His tagged spinach OEC16 transit peptide (unmodified, S54C, M64F, and S54C, M64F = tag TP) as bait peptides (see text for details). The purified Tat complexes obtained were subjected to by BN-PAGE and Western detection using anti-TatB antibodies as described legend to Fig. 6. The additional high molecular mass signal (*) is found with all bait peptides carrying the M64F mutation.

| | protein name | sequence coverage % | alignment score | intensity | $log_{10}$(iBAQ) |
|---|---|---|---|---|---|
| 1 | tag TP | 85,8 | 323,3 | 5,6E+09 | 8,9 |
| 2 | unknown | 3,9 | -2,0 | 7,3E+09 | 8,5 |
| 3 | *Pisum sativum* TatC | 7,7 | 13,9 | 3,6E+08 | 7,5 |
| 4 | *Pisum sativum* TatB | 19,4 | 33,1 | 1,5E+08 | 7,1 |
| 5 | PsaN | 5,9 | 8,0 | 6,3E+07 | 7,0 |
| 6 | PsbS | 11,9 | 22,0 | 9,6E+07 | 6,9 |
| 7 | AP-3 complex subunit delta | 2,1 | 7,9 | 3,6E+08 | 6,9 |
| 8 | PsaF | 12,6 | 19,2 | 6,4E+07 | 6,8 |
| 9 | 30S ribosomal Rps7 | 6,5 | 7,0 | 4,6E+07 | 6,8 |
| 10 | Chl a-b binding protein | 7,1 | 19,7 | 4,5E+07 | 6,7 |
| 11 | Chl a-b binding protein | 6,7 | 15,2 | 5,2E+07 | 6,7 |
| 12 | PsaL | 9,8 | 15,8 | 2,8E+07 | 6,6 |
| 13 | scarecrow-like protein 23 | 2,9 | 8,9 | 1,6E+07 | 5,9 |
| 14 | LHC-like protein | 3,9 | 7,2 | 7,9E+06 | 5,7 |
| 15 | PetC (Rieske FeS protein) | 17,8 | 11,5 | 3,7E+06 | 5,6 |
| 16 | ATP-synthase subunit/AAA-domain | 4,7 | 14,1 | 1,5E+07 | 5,6 |
| 17 | coatomer zeta-3-like | 6,1 | -2,0 | 2,0E+06 | 5,6 |
| 18 | actin-like proteins | 7,4 | 12,8 | 8,0E+06 | 5,6 |
| 19 | G-patch-domain protein TGH | 1,2 | 6,6 | 2,0E+07 | 5,6 |
| 20 | Psb33-like protein | 4,4 | 7,8 | 5,1E+06 | 5,5 |
| 21 | PsbQ (OEC16) | 8,9 | 6,7 | 3,0E+06 | 5,4 |
| 22 | COBW domain-protein 1-like | 3,1 | 6,8 | 4,9E+06 | 5,3 |
| 23 | unknown | 2,5 | -2,0 | 4,5E+06 | 5,3 |
| 24 | NADPH-protochlorophyllid oxidoreductase | 3 | 7,1 | 4,6E+06 | 5,3 |
| 25 | $K^+$-efflux-antiporter 3, chloroplast | 1,6 | 6,4 | 1,9E+06 | 4,8 |

**Supplementary Figure S5: Proteins identified by LC-MS/MS in affinity-purified Tat complexes**

Pea thylakoids were incubated with recombinant tagTP, washed, and solubilized with Digitonin. The sample was subjected twice to metal ion affinity chromatography (IMAC). Pooled elution fractions were separated by SDS-PAGE and the gel lane was excised and cut into pieces. These were treated with trypsin, and analyzed by LC-MS/MS. The results for each gel piece were combined and the determined masses of the tryptic fragments were matched against the respective calculated masses of tagTP as well as TatB and TatC from *Pisum sativum* and the proteome of *Cicer arietinum* (chickpea), a close relative of *Pisum sativum*.
