## Supplementary figures and images for "The trimeric thylakoidal Tat receptor complex consists of a homo-oligomeric TatC core with associated TatB subunits"

### Graphical Abstract

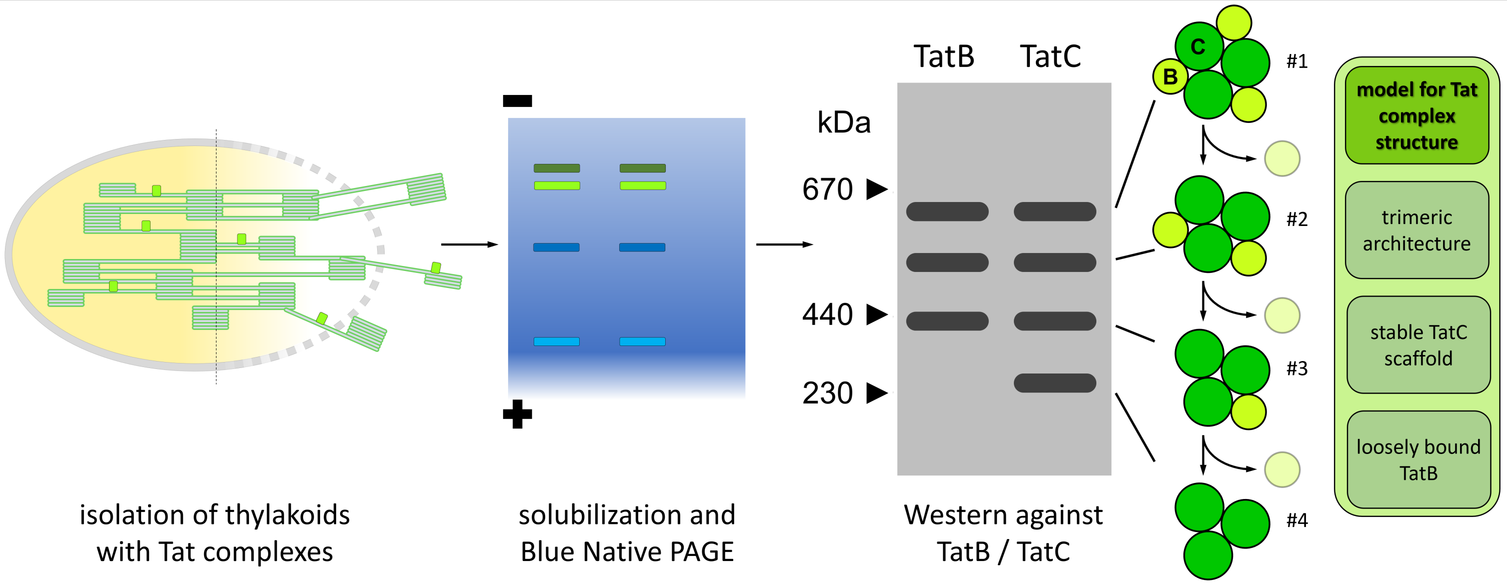
